# TrpA1 is a shear stress mechanosensing channel regulating intestinal homeostasis in Drosophila

**DOI:** 10.1101/2022.01.10.475734

**Authors:** Jiaxin Gong, Niraj K. Nirala, Jiazhang Chen, Fei Wang, Pengyu Gu, Qi Wen, Y. Tony Ip, Yang Xiang

**Affiliations:** Department of Neurobiology, University of Massachusetts Medical School, Worcester, MA 01605, USA; Program in Molecular Medicine, University of Massachusetts Medical School, Worcester, MA 01605, USA; Department of Physics, Worcester Polytechnic Institute, Worcester, MA 01609, USA

**Keywords:** calcium channel, *Drosophila*, enteroendocrine cell, intestine, mechanosensing, stem cell, TrpA1

## Abstract

Adult stem cells are essential for maintaining normal tissue homeostasis and supporting tissue repair. Although genetic and biochemical programs controlling adult stem cell behavior have been extensively investigated, how mechanosensing regulates stem cells and tissue homeostasis is not well understood. Here, we show that shear stress can activate enteroendocrine cells, but not other gut epithelial cell types, to regulate intestine stem cell-mediated gut homeostasis. This shear stress sensing is mediated by transient receptor potential A1 (TrpA1), a Ca^2+^-permeable ion channel that expressed only in enteroendocrine cells among all gut epithelial cells. Genetic depletion of TrpA1 or modification of its shear stress sensing function causes reduced intestine stem cell proliferation and intestine growth. We further show that among the TrpA1 splice variants, only select isoforms are activated by shear stress. Altogether, our results suggest the naturally occurring mechanical force such as fluid passing generated shear stress regulates intestinal stem cell-mediated tissue growth by activating enteroendocrine cells, and *Drosophila* TrpA1 as a new shear stress sensor.

## Introduction

Tissue maintenance involves resident stem cells that provide regenerative capacity to replenish lost cells due to aging or damage (1, 2). Intestine is one of the most active regenerative tissues in the human body because of a high rate of cell turnover (3, 4). In mammalian intestines, at least two populations of intestinal stem cells (ISCs) that have different proliferation rates serve to provide progenitors for routine tissue maintenance and damage-induced regeneration (5, 6). This multifaceted regenerative system allows rapid adaptive growth of the mammalian intestine in response to change of food intake or daily assault of ingested substances (3, 4). The regulation of ISC division therefore is essential to maintain the integrity of intestinal epithelia that is tightly linked to the health and survival of an individual (3, 7).

The midgut of adult fruit fly, *Drosophila melanogaster*, contains resident ISCs that serve as the major dividing cells for intestinal tissue homeostasis (3, 4, 8). Approximately a thousand ISCs are distributed individually and evenly along the midgut epithelium (9, 10). The midgut epithelium, similar to the mammalian intestine, is largely a single layer of mature enterocytes (ECs) that contain tight junctions to provide both absorptive and barrier functions. ISCs, precursor cells called enteroblasts (EBs), and enteroendocrine cells (EEs) are smaller cells that situated closer to the basal side of the midgut epithelium and serve to support various functions of the intestine (3, 4, 8). The coordination of ISC division, EB differentiation, and physiological functions of EEs and ECs requires many evolutionarily conserved pathways (2, 4, 8, 11). The investigation of these pathways in the adult *Drosophila* midgut as a model organism have provided important insights into intestinal regeneration and tissue wellness in mammalian systems (2, 4, 8, 11).

Regulation of ISC activity and intestinal homeostasis involves physiological signals such as growth factors, hormones, and innervation of the nervous system, as well as external signals from interaction with commensal and ingested microbes, and ingested food particles and fluids (2, 4, 8, 11). The intestinal epithelium is wrapped around by visceral muscle, and the rhythmic peristalsis propels ingested food particles and fluid down the gut lumen. While there is presumed mechanosensing processes, e.g., stretch and shear stress, caused by food particles and fluid passing through the intestinal lumen, the detailed mechanisms of mechanosensing in the gut epithelium and how that may regulate intestinal tissue homeostasis are just beginning to be discovered (12-16). Here we show that a member of the Ca^2+^-permeable transient receptor potential (Trp) channels family, *Drosophila* TrpA1, which has been previously known to be a polymodal channel for sensing heat and noxious chemicals (17-20), mediates shear stress sensing of enteroendocrine cells to regulate ISC activity.

## Results

### EEs are activated by shear stress *ex vivo*

Stretch, compression, and shear stress are physiological relevant mechanical forces associated with gut peristalsis and passing of food and fluid (Fig. 1A). Here, we determined how midgut epithelial cells respond to acute mechanical stimulation, in an *ex vivo* preparation after fresh dissecting adult *Drosophila* midgut. To stimulate epithelial cells with shear stress, we opened up the dissected gut to expose the lumen, placed the opened midgut in a custom-made microfluidic chamber, and delivered quantitative fluidic shear stress stimulation (Fig. 1B and S1A). The strengthen of shear stress was determined by adjusting the speed of saline flow driven by the pump (Fig. 1B). Cells in the normal environment of the gastrointestinal tract are estimated to be subjected to fluid shear stresses up to 30 dyne/cm^2^ (21). In this study, we stimulated adult midgut epithelium with 0.5 dyne/cm^2^, a small magnitude likely within the physiological range. Meanwhile, stretch was provided by pinning down the intact, non-opened midgut over a substrate and pushing down a section over a cavity (Fig. 1D and S1B). Such downward tissue movement increased the distance between the epithelial cells that were investigated by Ca^2+^ imaging in the region over the substrate (boxed area in Fig. 1D), indicating that stretch was successfully applied. On the other hand, compression was generated by direct pushing of opened midgut epithelium using a fire polished glass probe mounted on a piezo actuator (Fig. 1F and S1C). Mechanical responses are often associated with elevation of cytosolic Ca^2+^ signals (16, 22-26). As such, to measure responses of different midgut cell types, we expressed UAS-GCaMP6s, a genetically encoded Ca^2+^ sensor (27), under the control of various drivers, including Myo1A-Gal4 for ECs, Esg-Gal4 for ISCs/EBs, and TrpA1-D-2A-Gal4 for EEs (13, 16, 18, 28, 29). TrpA1-D-2A-Gal4, generated by inserting 2A-Gal4 to the C-terminus of TrpA1 genomic locus in the knockin allele that expressed only TrpA1-D isoform (18), was expressed in the midgut specifically in EEs (see below). By time-lapse imaging for monitoring GCaMP responses, we found that EEs, but not ECs or ISCs/EBs, quickly responded to shear stress stimulation (Fig. 1B-1C, and 1G-1I), indicating that EEs among gut epithelial cells are specifically activated by shear stress mechanical stimulation. Further, EE responses to shear stress was also confirmed when UAS-GCaMP6s was directed by other EE drivers, including prospero-Gal4, tachykinin (TK)-2A-Gal4, DH31-Gal4, and TrpA1-C-2A-Gal4 (Fig. 1J) (18, 28, 30-32). On the other hand, stretch or compression did not activate Ca^2+^ responses in EEs (Fig. 1D-1G), ISCs/EBs (Fig. 1H), or ECs (Fig. 1I). Previous reports suggest the involvement of a small population of ISCs/pre-EEs that express Piezo, and of all EBs expressing the Misshapen-Yorkie pathway, to be responsive to mechanical stimulation *in vivo* (13, 16). Under the current experimental condition, however, we did not detect mechanical response of these cells. Possible explanations include the *ex vivo* experimental conditions and the sensitivity of the assays.

**Fig. 1.**
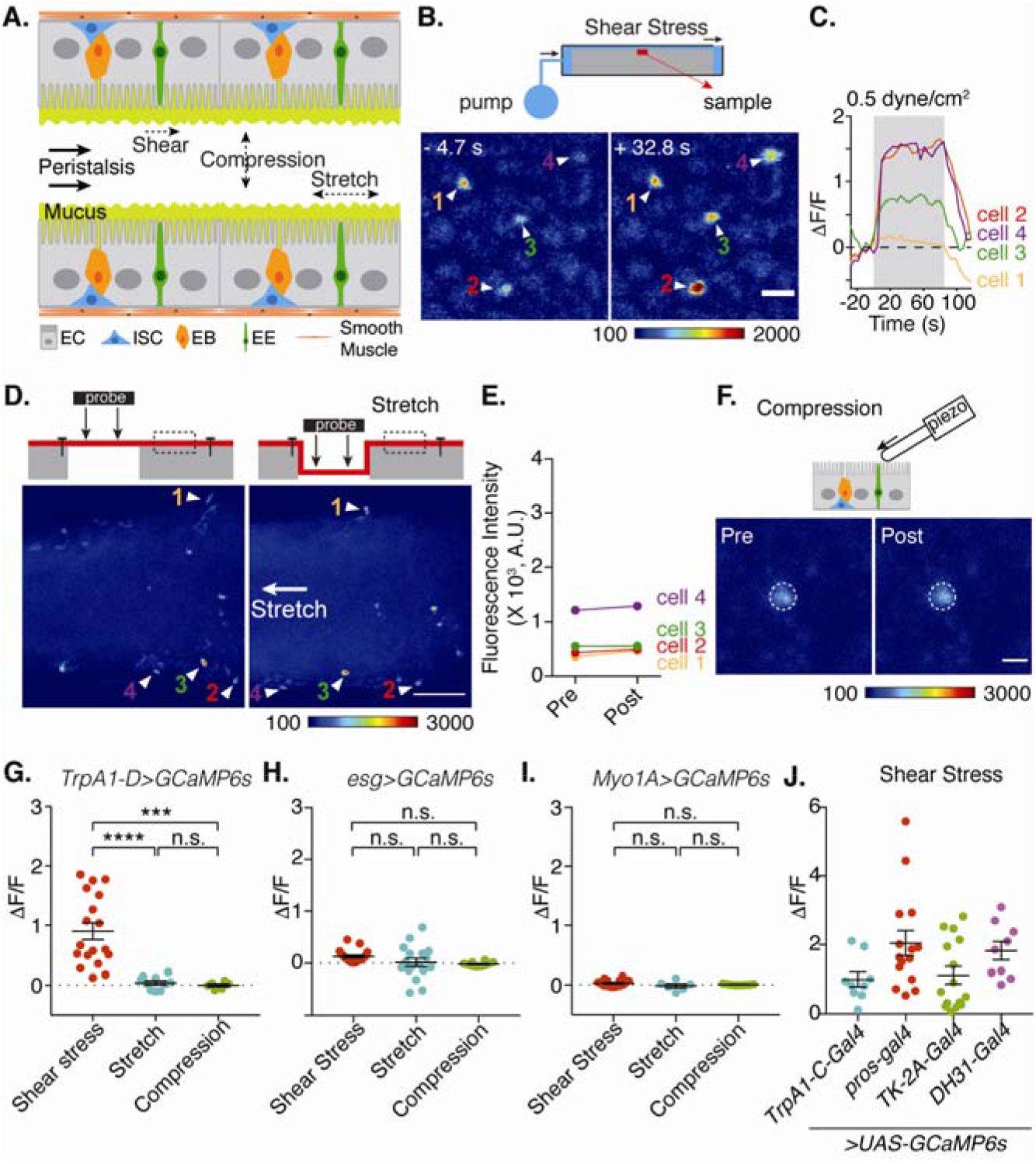
Shear stress activates adult *Drosophila* midgut EEs. (**A**) Schematics showing fly midgut cell types and three major forces: shear stress, compression and stretch, they would experience from peristalsis. (**B**) Ca^2+^ signals of EEs before (-ss) and during (+ss) shear stress stimulation. Scale bar, 10 μm. Top cartoon shows shear stress is delivered to gut lumen side in a microfluidic chamber. Flow rate is controlled by a syringe pump. All imaging was performed on posterior midgut (R4 or R5) of 20 days old adult female flies. (**C**) Ca^2+^ signal traces show slow adapting shear stress responses of four EEs (marked in panel B). (**D**) Ca^2+^ signals of EEs before (left subpanel) and after (right subpanel) stretch stimulation. Top cartoon shows the stretch assay, in which gut is fixed at two ends and deflecting gut towards hollow space by 500 μm with a probe above would cause ∼80% extension of the gut. Arrowheads and numbers denote representative cells before and after stretch. Scale bar, 25 µμm. Arrow denotes the direction of stretch. (**E**) Absolute GCaMP signals before and after stretch for the cells marked in panel D. (**F**) Representative GCaMP signals of EE toward 10 µm compression stimulation. Top cartoon shows the compression assay. Cells are stimulated from the lumen side by a blunt probe (about ∼10 µm in diameter for the tip) mounted on a piezo actuator. Scale bar, 5 µm. (**G**-**I**) Quantification summary of GCaMP responses of the midgut cells to shear stress, stretch or compression forces stimulations. In panel G, EEs are labelled by *TrpA1-D-Gal4/UAS-GCaMP;* in panel H, ISC/EBs are labelled by *esg-Gal4*; in panel I, ECs are labelled by *Myo1A-Gal4* (**I**). n >= 6. One-way ANOVA with Tukey’s multiple comparisons test. ^***^*p* < 0.001;^****^*p* < 0.0001. (**J**) Shear stress responses of EEs labelled by 4 other independent drivers. n >= 9. One-way ANOVA with Dunn’s multiple comparisons test. All showed significant response, and no significant differences were found among the four groups.

### TrpA1 has extensive expression in EEs

Among the over 64 Ca^2+^ channels encoded by the *Drosophila* genome, only very few have significant expression in the adult midgut (flybase). The stretch sensitive channel Piezo was previously shown to be expressed in a small number of ISCs/pre-EEs (16). TrpA1 is another Ca^2+^ channel previously shown to be expressed in EEs (19), while another report suggested TrpA1 expression is in ISCs/EBs (20). To further analyze the expression pattern of TrpA1, we utilized our recently generated TrpA1 knockin alleles, resulting in direct fusion of either GFP or 2A-Gal4 to the C-terminus of endogenously expressed TrpA1 protein (18). Using an available TrpA1 antibody (33), which may have a relative low titer, we have previously validated the pattern of expression is largely consistent between endogenous TrpA1 proteins and these knockin alleles in neurons (18). Immunostaining against GFP in TrpA1-GFP knockin flies revealed the only detectable signal of TrpA1 in adult midguts were in EEs, as evidenced by the co-staining with the pan-EE marker Prospero (Pros) (Fig. 2A). The Pros protein staining by the monoclonal antibody used is highly specific in probably all EEs in adult and larval midguts (9, 10). We examined carefully for expression of TrpA1 in the whole adult midgut and concluded that the only detectable expression is in EEs and any expression in other cell types is beyond our detection limit. To validate that TrpA1 expressed in EEs is functional, we applied AITC, a specific agonist of TrpA1 to stimulate midgut epithelium *ex vivo*. GCaMP imaging revealed that EEs, marked by either TrpA1-D-2A-Gal4 or TK-2A-Gal4 (18, 30), robustly responded to AITC, whereas ISCs/EBs, driven by Esg-Gal4, failed to respond (Fig. 2B-2C). These results support the notion that EEs in adult midguts contain TrpA1 and therefore can respond directly to AITC stimulation.

**Fig. 2.**
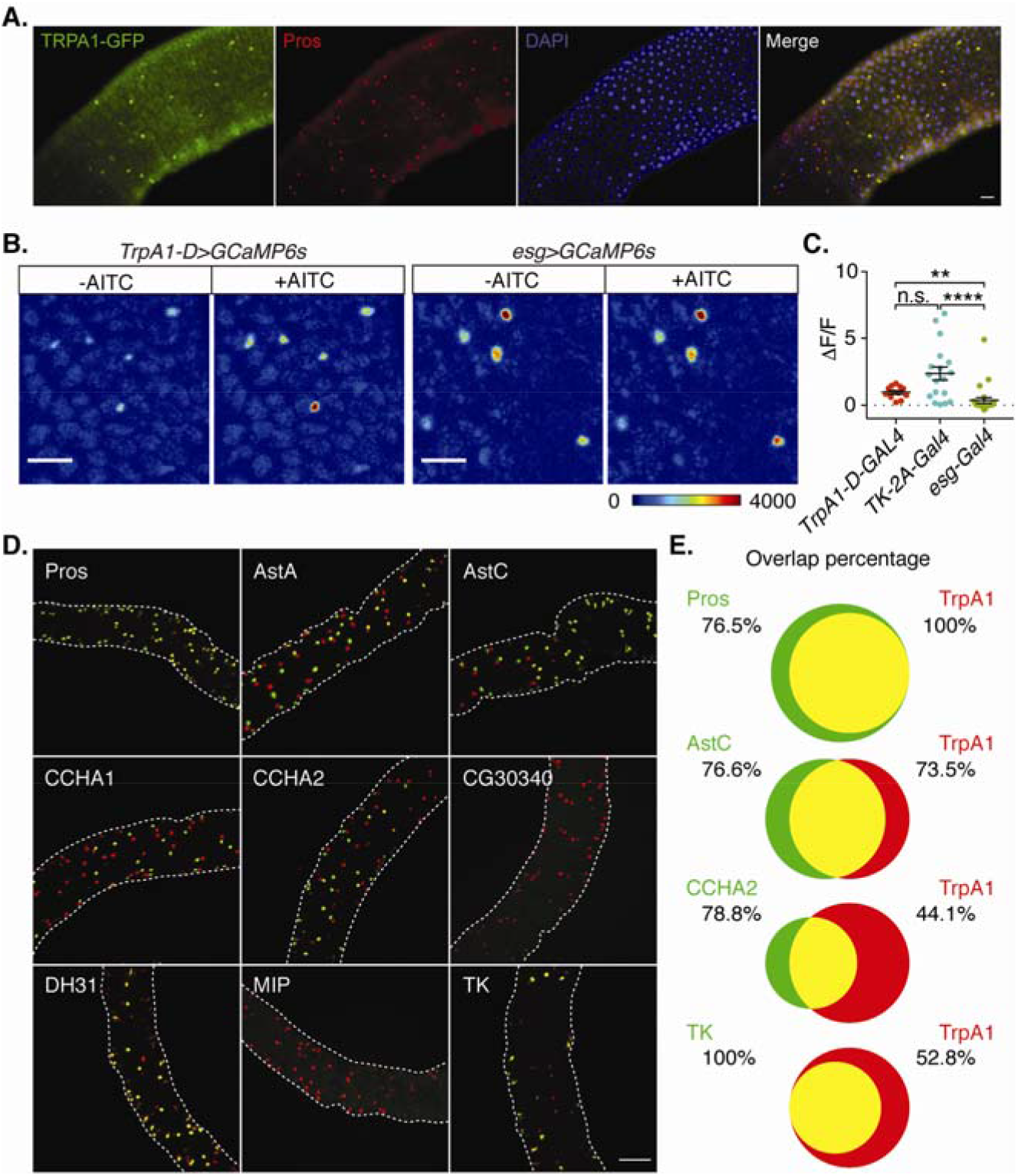
Expression of TrpA1 in midgut EEs. (**A**) co-staining of GFP and pan-EE marker Prospero (Pros) in *TrpA1-GFP* knock-in flies. Scale bar, 20 μm. (**B**) Representative images of Ca^2+^ responses to 100 mM AITC for EEs (*TrpA1-D>GCaMP6s*) and ISCs/EBs (*esg>GCaMP6s*) in posterior midgut. Scale bar, 25 µm. (**C**) Summary of Ca^2+^ responses to 100 mM AITC for EEs (labelled by *TrpA1-D-Gal4* or *TK-2A-Gal4*) and ISCs/EBs (labelled by *esg-Gal4*). n >= 12. One-way ANOVA with Dunn’s multiple comparisons test. ^**^*p* < 0.01; ^****^*p* < 0.0001. (**D**) TrpA1 and peptides are co-expressed in posterior EEs. Peptide-expressing EEs are labelled in green under the control of *peptide-2A-Gal4*, and TrpA1-expressing EEs are labelled in red under the control of *TrpA1-2A-LexA. Pros-Gal4* marks all EEs. (**E**) Overlap percentage between TrpA1 and Pros, AstC, CCHA2, or TK. The number specifies the percentage of cells expressing the gene that also expresses the other gene in same group.

Different populations of EEs express various combinations of over 30 different peptide hormones (31, 34-36). These different populations of EEs may serve different physiological functions. By using available Gal4 driver lines of some of these hormone genes, we quantify the co-expression patterns of TrpA1 with various hormones. The results show extensive but variable overlap with multiple hormones. Assuming all EEs are Pros-positive, TrpA1 is expressed in approximately 77% of all EEs. While many hormones have variable overlap with TrpA1, all Tachykinin (TK) expressing EEs are also TrpA1-positive (Fig. 2D-2E). These results suggest that TrpA1 is expressed in a major population of EEs and overlap to a different degree with various gut hormone expression patterns, which may allow TrpA1 to regulate different physiological functions of the midgut.

### EEs shear stress sensing depends on TrpA1

The TrpA1 expression pattern suggests a possible role of TrpA1 in shear stress sensing of EEs. To test this possibility, we expressed UAS-TrpA1 RNAi, driven by the TrpA1-2A-Gal4 driver. TrpA1 knockdown led to abolishment of EE responses to shear stress in *ex vivo* preparations (Fig. 3A), suggesting that TrpA1 is responsible for shear stress sensing of EEs to increase intracellular Ca^2+^. To corroborate this result, we used a null TrpA1 mutant previously generated in our laboratory that deletes the entire TrpA1 (18). We found shear stress induced Ca^2+^ responses in EEs were also abolished in this TrpA1-KO allele (Fig. 3B). Alternative splicing produced 5 TrpA1 isoforms (Fig. S2A). Among them, four isoforms (TrpA1-A, B, C and D) are functional, whereas the fifth isoform TrpA1-E exhibited no function (18). Analysis of expression of individual TrpA1 isoforms using the isoform-specific knockin alleles revealed that all 4 functional isoforms were expressed in midgut (Fig. S2B) (18). When expressed in EEs under the TK promoter-controlled R61H07-Gal4, we found that TrpA1-D, but not TrpA1-A, rescued the shear stress phenotype of TrpA1-KO flies (Fig. 3B). The E1017K point mutant version of TrpA1-D, which has the channel function abolished (see below), did not show such rescue (Fig. 3B). On the other hand, both TrpA1-D and TrpA1-A rescued defective EEs responses to AITC of TrpA1-KO flies (Fig. 3C). These results demonstrate that TrpA1-A is still a functional channel in this transgenic rescue assay, and TrpA1-D but not TrpA1-A plays a role in EE shear stress sensing.

**Fig. 3.**
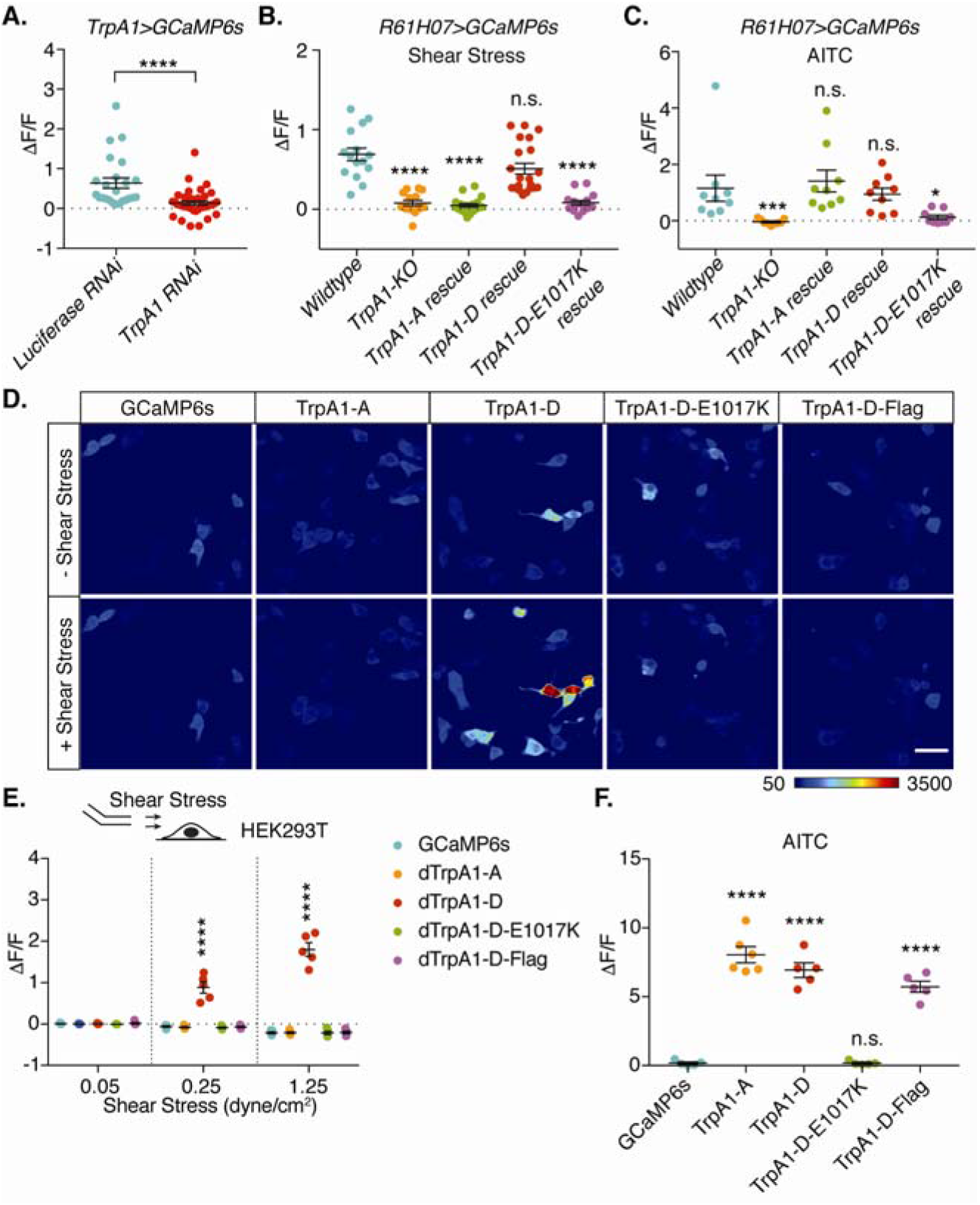
TrpA1 is a shear stress sensor mediating EE responses. (**A**) RNAi Knockdown of TrpA1 suppressed EEs shear stress responses. Luciferase RNAi served as control. N >= 24. Two-tailed Mann-Whitney test. ^****^*p*<0.0001. (**B**) Summary of EE responses to 0.5 dyne/cm^2^ shear stress in flies with genotypes indicated in figure. *R61H07-Gal4* was used to label EEs and drive expression of different TrpA1 isoforms. n >= 15. Comparison was made between wildtype and other groups using one-way ANOVA with Dunn’s multiple comparisons test. ^****^*p* < 0.0001. n.s., no significance. (**C**) Summary of EE responses to 100 mM AITC in flies with genotypes same as panel B. n = 9 for each group. Comparison was made between wildtype and other groups using one-way ANOVA with Dunn’s multiple comparisons test. ^*^*p* < 0.05; ^***^*p* < 0.001. n.s., no significance. (**D**) Representative images showing baseline (-shear stress) and peak (+ shear stress) responses to 1.25 dyne/cm^2^ shear stress, of HEK293T cells expressing GCaMP6s alone or together with Drosophila TrpA1-A, TrpA1-D, TrpA1-D-E1017K or TrpA1-D-Flag. Shear stress was delivered through a micropipette positioned 1 mm away from the center of imaging field. Scale bar, 50 µm. (**E**) Summary of HEK293T cell responses to stepwise shear stress stimulation. n = 5 for each group. Comparison was made between GCaMP6s alone group and other groups using two-way ANOVA with Dunnett’s multiple comparisons test. ^****^*p* < 0.0001. (**F**) Summary of HEK293T cell responses to 100 mM AITC. n >= 5 for each group. Comparison was made between GCaMP6s only group and other groups using two-way ANOVA with Dunnett’s multiple comparisons test. ^****^*p* < 0.0001. n.s., no significance.

### Select TrpA1 isoform is a shear stress sensor

To directly demonstrate TrpA1 as a mechanosensitive ion channel mediating response to shear stress, we utilized heterologous cells to examine whether TrpA1 could confer shear stress responses. We stimulated HEK293 cells transfected with *Drosophila* TrpA1 with shear stress. Cells transfected with TrpA1-D exhibited robust Ca^2+^ responses to shear stress, whereas cells transfected with TrpA1-A or GCaMP6s alone did not respond to shear stress (Fig. 3D-3E). These results indicate that select TrpA1 isoform, i.e., TrpA1-D but not TrpA1-A, is a shear stress sensor, and provide an explanation of why expression of TrpA1-D but not TrpA1-A rescued TrpA1-KO phenotype of shear stress sensing (Fig. 3B). Moreover, HEK293 cells expressing TrpA1-D bearing a point mutation in the putative pore domain (E1017K) again failed to respond to shear stress (Fig. 3D-3E), indicating ion conduction through TrpA1 is essential for the observed shear stress responses. Moreover, HEK293 cells transfected with either TrpA1-D or TrpA1-A, but not mutated TrpA1-D bearing the pore mutation, are activated by AITC (Fig. 3F), indicating that both TrpA1-D and TrpA1-A are AITC sensors.

### TrpA1 regulates ISC-mediated intestinal growth in adult midgut

The adult midgut epithelium is essentially a monolayer of ECs occupying most of the epithelium and the smaller-sized precursor cells and EEs located closer to the basement membrane. The adult midgut epithelium continues to grow after eclosion from pupal stage and the growth is largely supported by the division of ISCs. As flies age, the midgut epithelium accumulates more cells. This is exemplified by staining of β-Catenin that reveals the cell membrane, together with staining of Pros that marks EE nuclei (Fig. S3), with more cells are packed together in older fly guts. We next examined phenotypic changes in the adult midgut in TrpA1-KO flies, and found that when compared to control *w*^*1118*^ flies, TrpA1-KO flies exhibited severe defects in gut growth, although overall viability was unaffected. In TrpA1 mutants, the gut epithelium appeared morphologically normal, but older mutant guts resemble young guts with fewer cells packed together. Quantification of mitotic activity by the phosphorylated Histone3 (p-H3) staining revealed a drastically reduced cell division especially in older TrpA1 mutant flies when compared to similar aged *w*^*1118*^ control flies (Fig. 4A-4B). Most of the p-H3 staining in adult midguts likely represent ISC division, while EE division is also possible in older guts but should represent less than 5% of p-H3 counts (34). This reduced mitotic count corroborates the reduced cell numbers of all epithelial cell types including ECs, ISCs/EBs, and EEs (Fig. 4C-4E), further supporting the idea that reduction of ISC division after loss of TrpA1 leads to reduced overall cell numbers and therefore reduced growth in the adult midgut. In addition to midgut, TrpA1 is expressed in neurons (18). To manipulate TrpA1 in EEs, we combined prospero-Gal4 with elav-Gal80. In the resulting driver line, Gal80, the Gal4 repressor (37), is expressed in all neurons under the control of pan-neuronal promoter elav. When this driver was crossed to UAS-TrpA1-RNAi to knock down TrpA1 specifically in EEs, we found p-H3 counts were also markedly reduced to a degree similar to TrpA1-KO flies (Fig. 4F). These results indicate that TrpA1 functions in EEs to regulate ISC-mediated intestine growth.

**Fig. 4.**
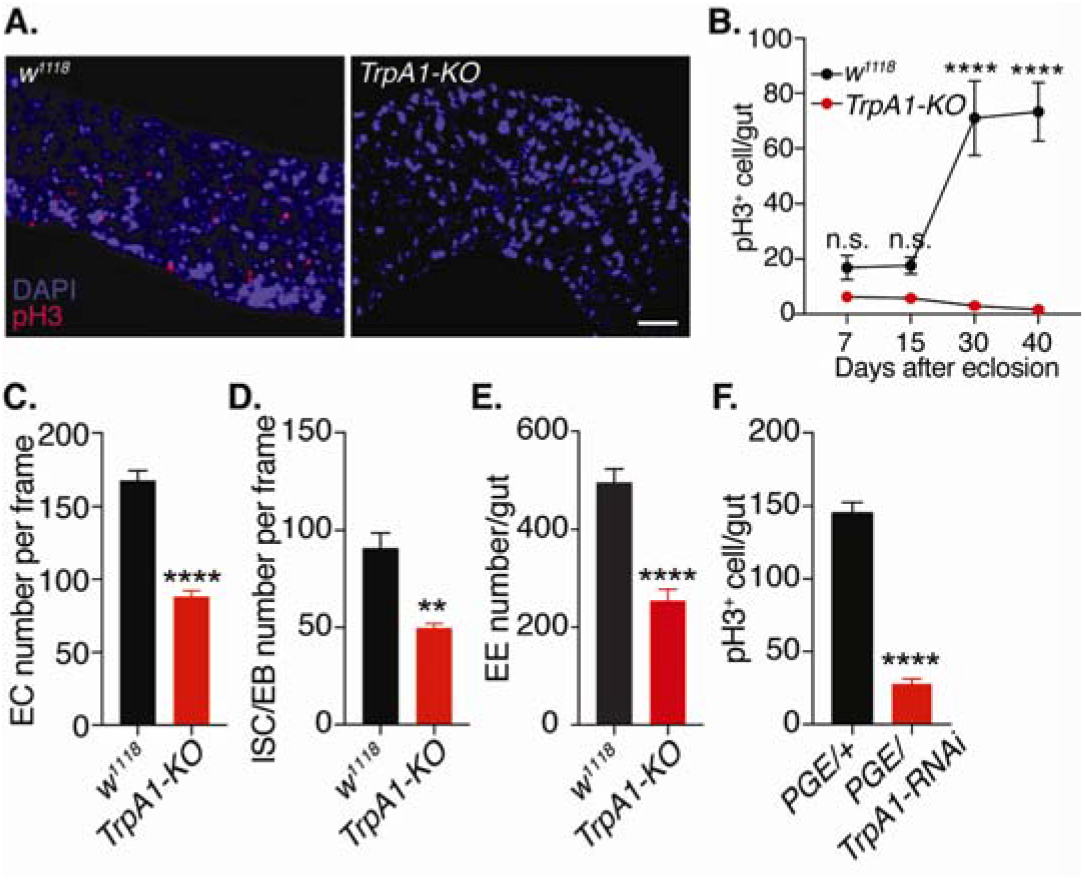
*TrpA1*-*KO* flies exhibited significant gut homeostasis defects. (**A**) Representative images of p-H3 staining (red) from 30-day old *w*^*1118*^ and *TrpA1*-*KO* flies. Scale bar, 20 μm. (**B**) p-H3 counts from whole gut at different ages. N >= 10 for each group. Two-way ANOVA with Bonferroni’s multiple comparisons test. ^****^*p* < 0.0001. n.s., no significance. (**C**-**E**) Number of ECs (**C**), ISCs/EBs (**D**) and EEs (**E**) in 40-day old *w*^*1118*^ and *TrpA1*-*KO* flies. n >= 6. Two-tailed Student’s t-test. ^**^*p* < 0.01; ^****^*p* < 0.0001. (**F**) RNAi knockdown of TrpA1 in EEs reduced ISCs proliferation. *Pros-Gal4, elav-Gal80* (*PGE*) was used to drive expression of TrpA1 RNAi only in EEs but not neurons. n>=11. Two-tailed Student’s t-test. ^****^*p* < 0.0001.

### TrpA1 shear stress sensing functions to control ISC proliferation

In addition to shear stress, *Drosophila* TrpA1 is also activated by heat, as well as noxious chemicals including electrophiles (e.g., AITC) and reactive oxygen species (ROS, e.g., H_2_O_2_) (18, 33, 38, 39). As we raised flies at the ambient temperature of approximately 23°C, which is below TrpA1 activation threshold, heat sensing functions of TrpA1 is unlikely to play a role. Because both electrophiles and ROS could be present in gut lumen, delineating the precise role of TrpA1-mediated mechanosensing requires a tool that can separate its shear stress sensitivity versus chemical sensitivity. We found that TrpA1-Flag knockin flies, which we have previously generated by inserting Flag to the C-terminus of TrpA1 immediately before the stop codon (18), satisfied this requirement. Specifically, we found that shear stress responses of EEs were abolished in TrpA1-Flag flies, similar to TrpA1-KO flies (Fig. 5A-5B). Intriguingly, unlike TrpA1-KO flies in which EE responses to irritant chemicals were also abolished, we found EEs in TrpA1-Flag flies retained their responses to H_2_O_2_ and AITC (Fig. 5C-5D). These results reveal TrpA1-Flag as a genetic tool to determine the *in vivo* function of EE shear stress sensing. Strikingly, we found that p-H3 counts were also markedly reduced in midguts of TrpA1-Flag flies (Fig. 5E), providing direct evidence that TrpA1 shear stress sensing functions can control ISC proliferation. Finally, we directly characterized sensory functions of TrpA1-Flag by expressing the construct in heterologous cells. Consistent with our findings in adult fly EEs, we found that HEK293 cells transfected with TrpA1-Flag failed to respond to shear stress (Fig. 3E), but exhibited normal responses to AITC (Fig. 3F). These results demonstrate that the shear stress mechanotransduction of TrpA1 has a specific structural requirement of the C-terminus that is distinguishable from those required for chemical nociception.

**Fig. 5.**
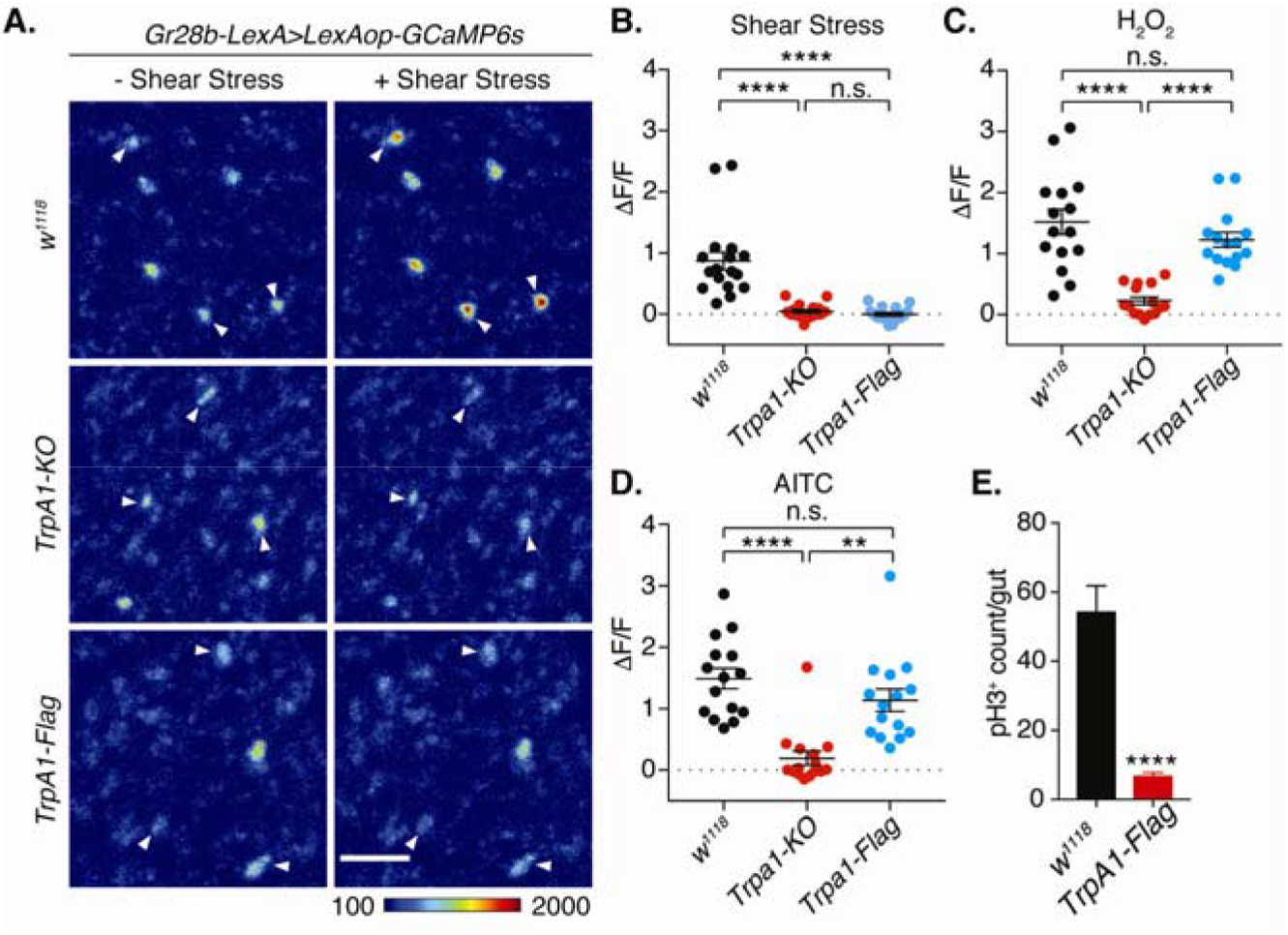
TrpA1 shear stress sensing function is essential for gut homeostasis. (**A**) GCaMP images of EEs before (-shear stress) and during (+ shear stress) 0.5 dyne/cm^2^ shear stress stimulation from *w*^*1118*^ control, *TrpA1-KO* and *TrpA1-Flag* flies. EEs are labelled by *Gr28b-LexA>LexAOP-GCaMP6s*. Arrowheads denotes representative EE cells. Scale bar, 20 μm. (**B**) Ca^2+^ responses of EEs to shear stress from *w*^*1118*^ control, *TrpA1-KO* and *TrpA1-Flag* flies. n >= 15. One-way ANOVA with Dunn’s multiple comparisons test. ^****^*p* < 0.0001. n.s., no significance. (**C**) Ca^2+^ responses of EEs to 10 mM H_2_O_2_ from *w*^*1118*^ control, *TrpA1-KO* and *TrpA1-Flag* flies. n = 15 for each group. One-way ANOVA with Tukey’s multiple comparisons test. ^****^*p* < 0.0001. n.s., no significance. (**D**) Ca^2+^ responses of EEs to 100 mM AITC from *w*^*1118*^ control, *TrpA1-KO* and *TrpA1-Flag* flies. n = 15 for each group. One-way ANOVA with Dunn’s multiple comparisons test. ^**^*p* < 0.01; ^****^*p* < 0.0001. n.s., no significance. (**E**) Mitotic p-H3^+^ cell count from whole gut of 40 days old *w*^*1118*^ control and *TrpA1-Flag* flies. n = 12 for each group. Unpaired Student’s t-test. ^****^*p* < 0.0001.

### TrpA1 modulates Ca^2+^ oscillation within ISCs that regulates proliferation

The expression of TrpA1 in EEs that leads to regulation of ISC division suggests a mechanism that involves communication between the two cell types. A previous report indicates that Ca^2+^ oscillation within ISCs regulates their division (29), although the mechanism that establish and regulates this ISC Ca^2+^ oscillation is not yet known. When we examined this spontaneous Ca^2+^ oscillation in ISC/EB cell nests, there was an almost complete loss of these signals in TrpA1-KO flies (Fig. 6A-6B). Moreover, Ca^2+^ oscillation within ISCs was also abolished in TrpA1-Flag flies (Fig. 6A-6B). These findings suggest that shear stress sensing of TrpA1 in EEs controls Ca^2+^ oscillation within ISCs, and therefore should play a role in the regulation of ISC proliferation.

**Fig. 6.**
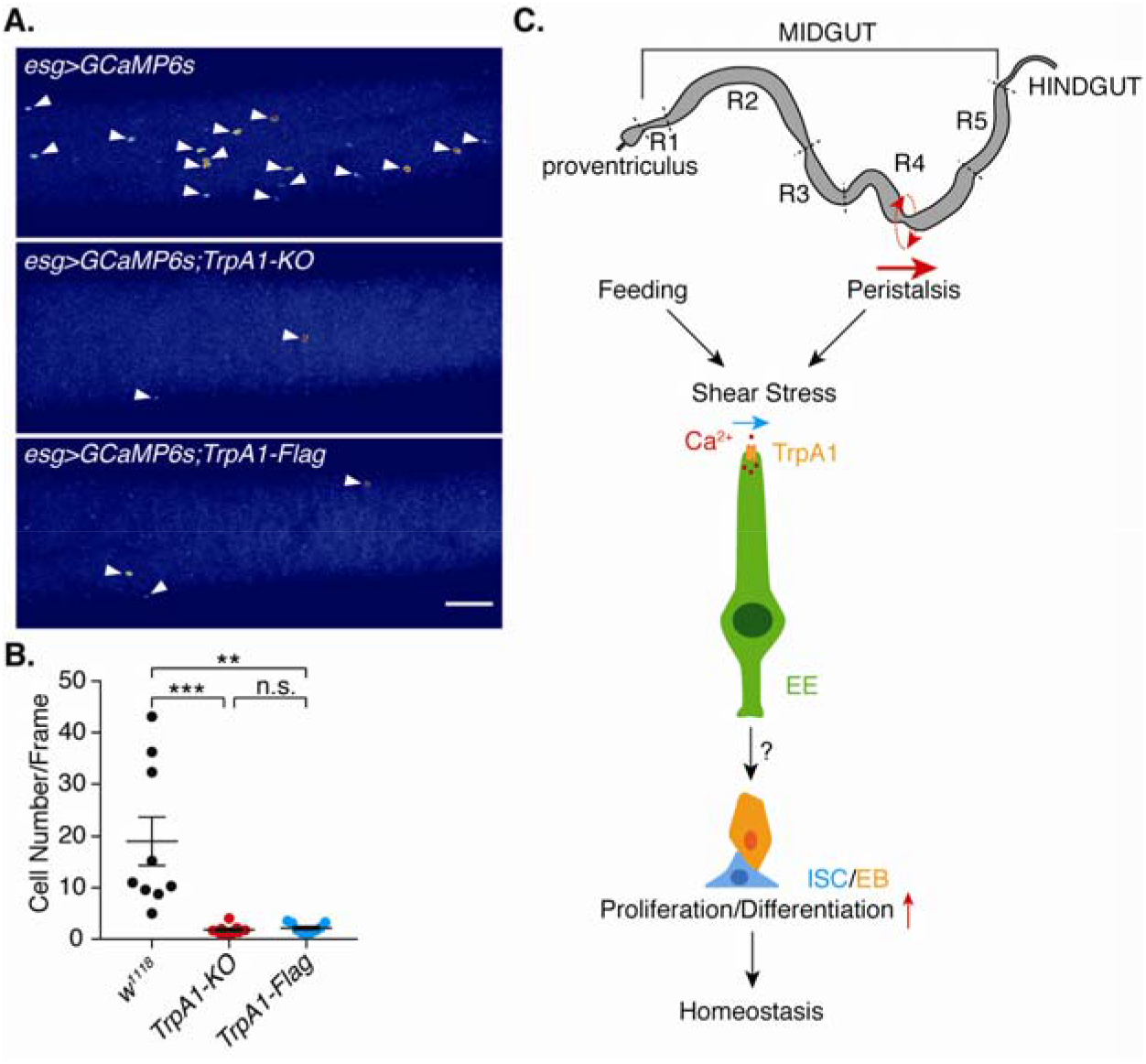
TrpA1 shear stress sensing regulates Ca^2+^ oscillation in ISCs. (**A**) Sample images of spontaneous Ca^2+^ oscillation in ISCs/EBs (*esg-Gal4>UAS-GCaMP6s*) from *w*^*1118*^ control, *TrpA1-KO* and *TrpA1-Flag* flies. Scale bar, 50 µm. (**B**) Oscillating cell count per frame during 10 min tracking from 20 days old *w*^*1118*^ control, *TrpA1-KO* and *TrpA1-Flag* flies. n >= 9. One-way ANOVA with Dunn’s multiple comparisons test. ^**^*p* < 0.01; ^***^*p* < 0.001.

## Discussion

In this report, we have uncovered a novel mechanism by which shear stress as a naturally occurring mechanical force can regulate stem cell division and tissue homeostasis in adult *Drosophila* midgut. In our model (Fig. 6C), shear stress sensing is mediated by EEs. Because a major function of EEs is to secret peptide hormones to regulate intestinal and whole-body physiology, our findings further suggest a possibility that shear stress induced Ca^2+^ elevation in EEs trigger peptide release to control ISCs division.

A previous report in *Drosophila* midgut indicates that mechanical stretch can activate the Piezo channel expressed in a subset of ISCs that will differentiate into EEs, to regulate ISC proliferation (16). This previous report, together with our study here, indicate that multiple types of mechanical forces act in parallel to orchestrate ISCs behavior. While stretch may directly regulate ISC proliferation by activating ISCs, shear stress indirectly controls ISC proliferation by activating EEs that subsequently signal to ISCs. Therefore, both cell-autonomous and cell-non-autonomous mechanisms are involved in mechanical regulation of ISC behavior.

We demonstrated in our study a novel function of TrpA1 as a shear stress mechanosensing Ca^2+^ channel. TrpA1 has been shown to be a polymodal nociceptor that transduce heat and noxious chemicals. Our results presented here have expanded the versatility of this Ca^2+^ channel in regulating tissue homeostasis. Not only that TrpA1 has this additional mechanosensory function, but we have further demonstrated that there is a specific structural involvement for shear stress that is separable from chemosensory functions of TrpA1. This structural requirement is selectively abolished when the small Flag sequence is attached to the C-terminus. The specific structural requirement of TrpA1 in mechanotransduction is further supported by the functional contribution of TrpA1-D but not TrpA1-A isoform. The sequence difference between these two splicing variants is ∼100 a.a. in the N-terminus. Meanwhile, the FLAG tag was inserted at the C-terminus. One speculation is that the 3D structure of the whole TrpA1 protein allows close proximity interaction of the N and C termini, both are facing the cytoplasm. While the detailed structural requirement is not known, our work provides a conceptual framework based on *in vivo* evidence to further analyze this evolutionarily conserved function of TrpA1 as a versatile ion channel to regulate mechanosensing and tissue homeostasis.

A previous study indicates that the Piezo channels mediate the shear stress responses of dissociated mouse EEs *in vitro* (25). We found that responses of *Drosophila* EEs to shear stress were unaffected in Piezo-KO flies (Fig. S4), indicating that TrpA1, instead of Piezo, is involved in shear stress sensing of *Drosophila* EEs.

Altogether, our studies provide new cellular and ion channel mechanisms of how mechanical forces regulate adult stem cell behavior and tissue growth. Because TrpA1 has been shown to express in mammalian gut with high levels in EEs (40). it will be of interest to determine whether TrpA1 could mediate shear stress sensing of EEs and hence play a conserved role in gut homeostasis in mammals.

## Acknowledgments

We thank the Perrimon lab for sharing reagents, members of Ip lab and Xiang lab for helpful discussion. Stocks were obtained from the Bloomington Drosophila Stock Center. Y.T.I. was supported by NIH grants (DK083450, GM107457). Y.T.I. is a member of the UMass Center for Clinical and Translational Science (UL1TR000161). Y.X. is supported by the National Institutes of Health under award number 1R21NS107924, the Human Frontier Science Program Young Investigator award RGY0090/2014, and the Martina Stern Memorial Fund.

---

**Fig. S1.**
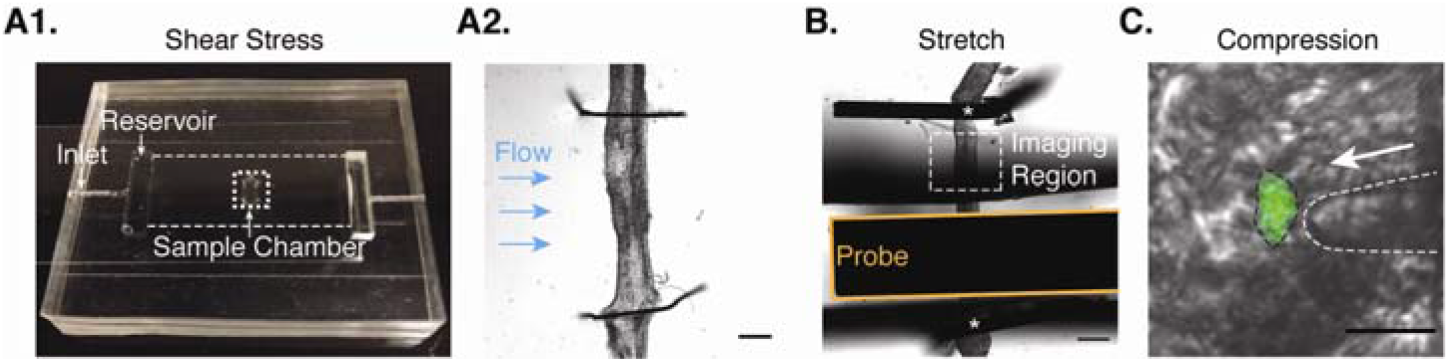
Force delivery paradigms. (**A1-A2**) Shear stress was delivered through a microfluidic chamber as in (**A1**). Channel dimension is 22 mm (L) X 12 mm (W) X 200 μm (H). The inlet connected to the syringe pump, and right reservoir (not labelled) was left open. The gut was opened to expose lumen and was placed in the sample chamber. Posterior midgut was cut open and fixed by fine tungsten wires as in (**A2**). Homogeneous laminar flow generates shear stress stimulation to samples. Scale bar, 200 μm. (**B**) Stretch stimulation paradigm. Intact (non-opened) gut was fixed at two ends (star marks) on a custom chamber over hollow space. Deflecting gut towards hollow space by a probe would stretch the gut. The imaging region (white dashed box) was marked. Scale bar, 200 μm. (**C**) Compression was achieved through direct pressing the cells of interest by an end fire polished glass probe which is mounted on a piezo actuator. Scale bar, 10 μm.

**Fig. S2.**
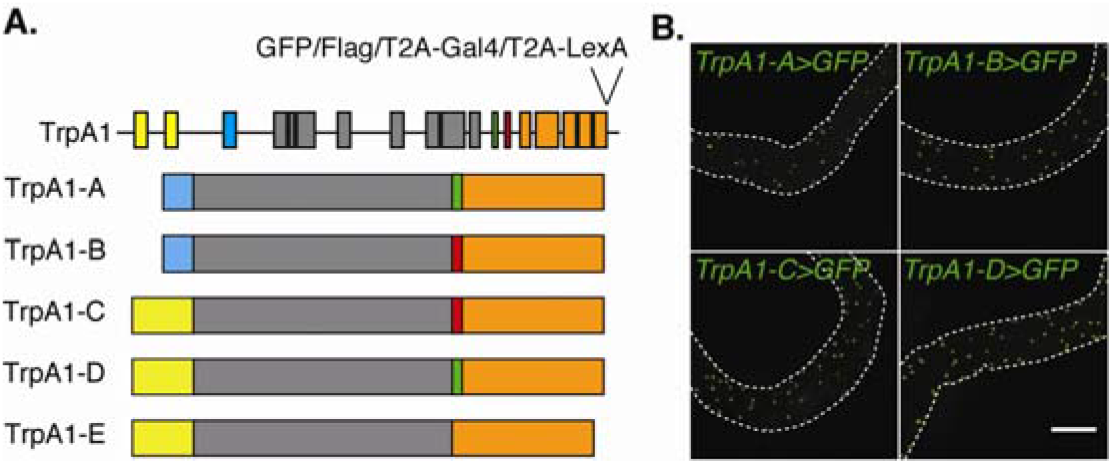
TrpA1 splicing isoforms and expression in adult gut. (**A**) TrpA1 alternative splicing gives rise to five isoforms that differ in the pre-ankyrin repeat N-terminus (blue or yellow), and in the linker domain (green or red) that connects ankyrin repeats to transmembrane domains. Genomic modification by isoEXPRESS generates TrpA1 isoform specific knockin flies bearing GFP, Flag, or T2A-Gal4 at the C terminus. See Gu et. al. 2019 for details. (**B**) Expression of individual TrpA1 isoform was detected in the midgut, using the isoform-specific 2A-Gal4 line driving UAS-6XGFP. Only TrpA1-A, B, C, D isoform expression in the R2 gut region was shown. Dashed lines denote the gut. Scale bar, 200 µm.

**Fig. S3.**
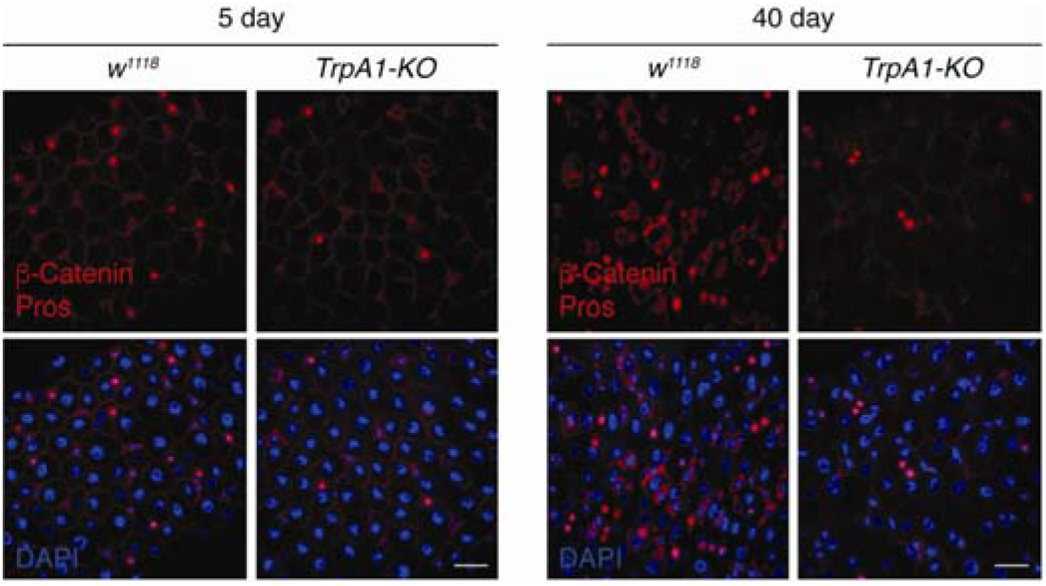
TrpA1 knockout causes gut homeostasis defects in aged flies. Immunostaining images of β-Catenin, which marks cell membrane, and Pros, which marks EE nuclei, of midguts from 5 days and 40 days old *w*^*1118*^ and *TrpA1-KO* flies. Scale bar, 20 µm.

**Fig. S4.**
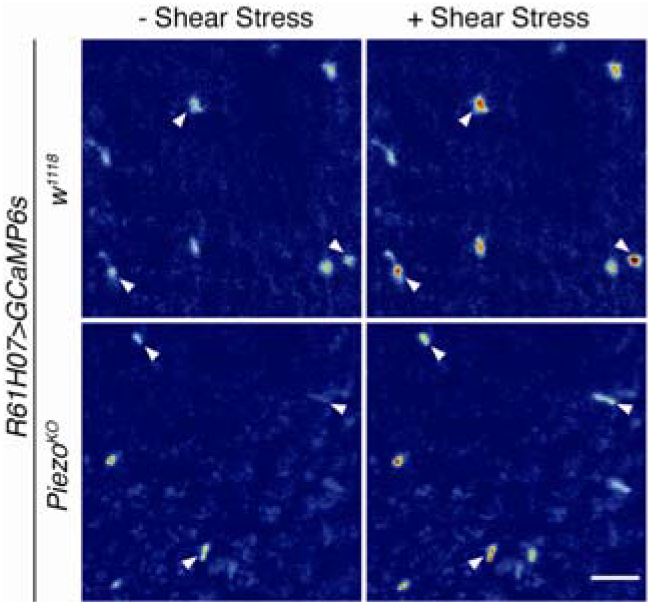
Shear stress responses of EEs was intact in Piezo knockout flies. Representative GCaMP signal images of EEs before (-shear stress) and during (+ shear stress) 0.5 dyne/cm^2^ shear stress stimulation from *w*^*1118*^ control and *piezo*^*KO*^ flies. Expression of GCaMP6s in EEs was driven by *R61H07-Gal4*. Arrowheads denotes responsive EE cells. Scale bar, 20 µm.

## Materials and Methods

### *Drosophila* stocks and transgenic lines

All fly stocks were raised on low yeast brown food, which was prepared by the University of Massachusetts Medical School *Drosophila* Resource Facility. Flies were maintained at room temperature unless otherwise noted. For GCaMP imaging studies, 20 days old adult female flies were used. For immunostaining studies, adult female flies were used at indicated age. The following fly strains were used in this study. *w*^*1118*^ (gift from Patrick Emery), *TrpA1-D-2A-Gal4* (pengyu), *esg-Gal4* (BL#84324), *myo1A-Gal4* (28), *UAS-GCaMP6s* (BL#42746, BL#42749), *TrpA1-C-2A-Gal4* (pengyu), *pros-Gal4* (BL#84276), *TK-2A-Gal4* (BL#84693), *DH31-Gal4* (BL#51988), *TrpA1-GFP* (pengyu), *AstA-2A-Gal4* (BL#84593), *AstC-2A-Gal4* (BL#84595), *CCHA1-2A-Gal4* (BL#84361), *CCHA2-2A-Gal4* (BL#84602), *CG30340-2A-Gal4* (BL#84611), *DH31-2A-Gal4* (BL#84623), *MIP-2A-Gal4* (BL#84651), *UAS-6XGFP* (BL#52262), *LexAop-6XmCherry* (BL#52267), *TrpA1-2A-Gal4* (18), *TrpA1-2A-LexA* (18), *UAS-luciferase-RNAi* (), *UAS-TrpA1-RNAi* (BL#36780), *TrpA1-KO* (18), *R61H07-Gal4* (BL#39282), *UAS-TrpA1-A* (18), *UAS-TrpA1-D* (18), *UAS-TrpA1-D-E1017K* (this work), *TrpA1-Flag* (18), *Gr28b-LexA* (lab stock), *LexAOP-GCaMP6s* (BL#44589).

### Mechanical shear stress assay in microfluidic chamber

The microfluidic chambers (See dimensions in Fig. S1) were fabricated using the standard soft lithography method. Polydimethylsiloxane (PDMS) solution (Sylgard 184, Dow Corning, Midland, MI) was poured over a customized channel mold and cured at 70°C for 2 h to produce a negative replica of the channels. The inlet of the microfluidic chamber was connected to a syringe pump (Harvard instrument), which controlled the flow rate. The shear stress magnitude of laminar flow in the chamber was calculated using the computational fluid dynamics (CFD) module in COMSOL Multiphysics 5.5. Results suggest that shear stress was uniform on the bottom surface of the microfluidic chamber with the magnitude of shear stress (*σ*) proportional to the flow rate (*Q*) as: 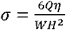, where *η* is the dynamic viscosity of the saline, and W and H are the width and height of the chamber, respectively (41).

The genetically encoded Ca^2+^ indicator GCaMP6s was used to determine responses of different gut cells. Briefly, the whole gut was dissected from 20 days old female flies in adult hemolymph like saline, which contains 108 mM NaCl, 5 mM KCl, 2 mM CaCl_2_, 8.2 mM MgCl_2_, 4 mM NaHCO_3_, 1 mM NaH_2_PO_4_, 5 mM trehalose, 10 mM sucrose, 5 Mm HEPES with pH adjusted to 7.4. Then the gut was slightly stretched and pinned down in the central sample chamber of the microfluidic device. Next, the posterior midgut was cut open with two sharp syringe tips and the internal content was removed to expose the lumen side of gut. Two “⌒” shaped tungsten wires (0.024 mm ID) were further used to fix and flatten the exposed region. The chamber was sealed with a 22 × 50 mm coverglass (Fisher Brand) and filled with AHL for further imaging studies. Ca^2+^ responses in gut cells were imaged as a time-lapse of 13 Z-stack at step of 2.5 □m every ∼5 s. Ca^2+^ signals from three to five cells for each gut with strongest responses were chosen for analysis in Image J. As ISC/EBs and a few EEs show robust Ca^2+^ oscillations, we avoid to analyze such cells based on the Ca^2+^ signals before stimulations.

### Mechanical stretch assay

To deliver tissue stretch stimulation to gut, a whole gut from 20 days old female fly was dissected in AHL and pinned down over a cavity (∼650 μm in width, Fig. S1). A probe made from a staple pin (with flat sides) with a width of ∼500 μm was used to press down the gut over the cavity, which caused stretch of the gut. Displacement of the glass probe was controlled by the programmable micromanipulator (MP285, Sutter instruments) at a speed of 6 mm/s. The gut was depressed 500 μm and stretched ∼80% based on ideal calculation (∼30% failure rate because of gut breakage from stretch). Imaging was focused on the region of posterior midgut at one shoulder of the cavity (dashed region in Fig. S1). Ca^2+^ responses in gut cells were imaged as a time-lapse of 15 Z-stack at step of 2.5 μm every ∼6 s. As ECs exhibit low Ca^2+^ signals, it’s difficult to identify the same cells before and after stretch. So we randomly picked one EC before and after stretch stimulations for analysis.

### Mechanical compression assay

Gut from 20 days old female fly was dissected and exposed as described in the shear stress assay. To mimic the compression stimulation, we used an end fire polished glass probe (5-10 μm in diameter) to compress the gut cells of interest (Fig. S1.). The glass probe was mounted on a piezo actuator (Thorlabs), which is driven by a stimulator (S88, The Grass Instrument). The compressive force was delivered at an angel of about 25 degree relative to the horizontal plane. Cells were compressed by a 10 μm displacement and then immediately released. Ca^2+^ responses in gut cells were imaged as a time-lapse of 10 Z-stack at step of 2.5 μm every 2 s.

### Cell culture

HEK293T cells were cultured in DMEM (Corning) supplemented with 10% fetal bovine serum (Sigma), 2 mM L-glutamine (Gibco), 1 mM sodium pyruvate (Gibco), 100 Unit/ml penicillin and 100 μg/ml streptomycin (Corning) at 37°C and 5% CO_2_. Cells were grown on 22 mm square coverglass coated with 0.2% gelatin. Transfection was done with 3 μg polyethylenimine (Polysciences) per 3.5 cm dish 1 day after plating. PcDNA3.1-TrpA1-A, pcDNA3.1-TrpA1-D, pcDNA3.1-TrpA1-D-E1017K, or pcDNA3.1-TrpA1-D-Flag were co-transfected with pcDNA3.1-GCaMP6s and pcDNA3.1 empty vector into cells at a ratio of 0.05:0.4:0.55 (in μg) per 3.5 cm dish. GCaMP imaging were performed 2 days after transfection. Cells were allowed to habituate for 15 mins at room temperature before experiments. Extra care was taken in handling the plates in order to minimize stimulation of the cells from shaking.

### Live Ca^2+^ imaging in HEK293T cells

The genetically encoded Ca^2+^ indicator GCaMP6s was used to determine responses of HEK293T cells. Imaging was conducted under a Zeiss LSM700 upright confocal microscope with a 20X/1.0 NA water immersion objective. External saline contained 130 mM NaCl, 3 mM KCl, 0.6 mM MgCl_2_, 1 mM CaCl_2_, 1.2 mM NaHCO_3_, 10 mM Glucose, 10 mM HEPES with pH adjusted to 7.3. Shear stress stimulations were delivered horizontally through a glass micropipette (OD, 1.2 mm; ID, 0.69 mm; Sutter Instrument) at 1 mm away from the center of the imaging field. Flow rate is controlled by a syringe pump (PHD 2000, Harvard Instruments), and shear stress response was tested at three levels (0.05, 0.25, 1.25 dyne/cm^2^) with a stepwise manner. Shear stress was calculated as 4*η*Q/(πr^3^) (42), where 1 is the viscosity of saline (the viscosity of water at room temperature, at a value of 1 mPa.s, was used), Q is the flow rate, π=3.14, and r is the internal radius of the micropipette (345 μm). Time-lapse tracking of GCaMP6s signals were performed at 2.5 frames/s and the mean value of GCaMP6s signals for each frame was quantified using ImageJ. Ca^2+^ responses under different shear stress levels were defined as the maximum fluorescence fold change (ΔF/F0 =(F-F0)/F0), where F is the maximum Ca^2+^ signal value for each level and F0 is the baseline before stimulation (without shear stress stimulation).

### Ca^2+^ oscillations in ISC/EBs

For imaging the spontaneous Ca^2+^ oscillations in ISC/EBs, GCaMP6s were expressed in ISC/EBs with esg-Gal4 to monitor the cytosolic Ca^2+^ changes. Imaging was focused on posterior midgut from 20 days old female flies. Adult hemolymph like saline, which contains 108 mM NaCl, 5 mM KCl, 2 mM CaCl_2_, 8.2 mM MgCl_2_, 4 mM NaHCO_3_, 1 mM NaH_2_PO_4_, 5 mM trehalose, 10 mM sucrose, 5 mM HEPES with pH adjusted to 7.4, was used for imaging. To reduce peristaltic movements of gut, 10 LJg/ml isradipine was added to imaging saline (heinrich jasper). Ca^2+^ signals were tracked as a time-lapse of 17-slice Z-stack at a step of 4 µm every ∼13.5 s. Every gut will be recorded 10 min. The number of cells showing oscillations and captured in each stack was counted (after maximal intensity projection) manually.

### Immunostaining

Midguts from female W1118, TrpA1-KI-Flag and TrpA1 mutant (different days as specified in legend) were fixed in 4% Formaldehyde (Polysciences) in 1XPBS for 2 hours under room temperature. The following rinses, washes, and incubations with primary and secondary antibodies were in the 1X PBS containing 0.5% BSA and 0.1% Triton-X100. The following primary antibodies were used: mouse anti-Pros (1:50 DSHB), mouse anti-armadillo (1:50 DSHB), rabbit anti-phospho-Histone3 (1:2000, Abcam), mouse Flag-M2 (1:500-sigma). Goat anti-mouse IgG conjugated to Alexa 568 (Invitrogen) and Goat anti-rabbit IgG conjugated to Alexa 555 (Invitrogen) were used as the secondary antibodies with 1:1000 dilution. The tissues were mounted in DAPI (antifade, Vectorshied, Vector Lab) and Images were captured with Nikon Spinning Disk confocal microscope and processed using ImageJ.

### Statistical Analysis

Statistical analysis was performed using Graphpad Prism 7 software. Error bars are standard errors of the mean (SEM). Three or more experiments using individual biological samples were performed for each of the experimental results presented in this report. The n in each figure represents the number of gut or cells counted in each experimental data point. For all statistics in this report, the error bar is standard error of the mean, and p value is from Student’s T-test: ^*^ is p<0.05, ^**^ is p<0.01, NS is no significance with p>0.05.

## Data and materials availability

all data is available in the manuscript or the supplementary materials.

